# Expression-based annotation identifies and enables quantification of small vault RNAs (svtRNAs) in human cells

**DOI:** 10.64898/2026.03.10.710617

**Authors:** Jake D. Sheppard, Pablo Smircich, María Ana Duhagon, Rafael Sebastián Fort

## Abstract

**Background:** Small non-coding RNAs (sncRNAs) play central roles in post-transcriptional gene regulation. In addition to canonical microRNAs (miRNAs), fragments derived from vault RNAs (vtRNAs), called small vault RNAs (svtRNAs), have been reported in human cells. However, the absence of a standardized annotation framework has hindered their systematic detection, quantification, and comparison across small RNA sequencing (small RNA-seq) studies.

**Methods:** We developed an expression-based annotation strategy to identify svtRNAs from human small RNA-seq datasets. Using FlaiMapper followed by structure and expression-based filtering, we generated two annotation sets: a stringent “miRNA-like” set enriched in Argonaute-associated datasets, and (ii) a broader “Total” set derived from total small RNA-seq libraries under relaxed structural constraints. We explored the expression of the annotated svtRNAs across the different datasets analyzed: multiple normal and tumor-derived human cell lines, including Argonaute immunoprecipitation datasets.

**Results:** We identified a repertoire of svtRNAs that are detected across independent datasets and, in several cases, reach abundance levels comparable to canonical miRNAs. Several highly abundant svtRNAs correspond to molecules with experimental validation from prior studies, supporting the robustness of our annotation strategy. Importantly, the same “dominant” (in terms of gene expression) svtRNAs emerged independently from Argonaute-associated and total small RNA datasets, supporting the idea of enzymatically consistent, reproducible svtRNA processing. We further identified svtRNAs derived from distinct vtRNA precursors that could share identical seed sequences, suggesting the possibility of svtRNA families with potential miRNA-like regulatory properties. We provide a standardized annotation that enables reproducible svtRNA quantification.

**Conclusions:** Our study establishes a comprehensive expression-based annotation resource for human svtRNAs. By enabling their systematic detection and reproducible quantification, we show that svtRNAs appear to represent an abundant component of the human small RNA landscape.

## Introduction

The central dogma of molecular biology proposed by Francis Crick in 1957, described the flow of information from genes to protein (Cobb 2017). Despite the early detection of non-coding RNAs (ncRNAs) in the human genome during the 1950s and 1960s, such as transfer RNA (tRNA), ribosomal RNA (rRNA), and small nuclear RNA (snRNA) (Micheel et al. 2021), it was not until the discovery and extensive investigation of microRNAs (miRNAs) in the 1990s and 2000s (see the discovery of lin-4 and let-7 in *Caenorhabditis elegans* as reviewed in (Aalto and Pasquinelli 2012); that the roles and impact of ncRNAs in the flow of genetic information began to be seriously examined. The development of DNA sequencing technologies, and in particular the establishment and expansion of small RNA sequencing (small RNA-seq) libraries, contributed to a marked increase in publications in the ncRNA field (Micheel et al. 2021). This led to annotation efforts such as the miRBase database (Griffiths-Jones 2006), which catalogs miRNAs across different species, including humans. Currently, 1,917 human miRNA precursors are annotated in miRBase (Kozomara et al. 2019), whereas more recent and conservative databases such as MirGeneDB compile a total of 514 human miRNA genes (Clarke et al. 2025). Despite the growing amount of small RNA sequencing data driven by decreasing sequencing costs, continued efforts are still required to annotate ncRNA biotypes in the human genome (Zerbino et al. 2020). In this context, beyond the classical small ncRNAs (miRNAs, piRNAs, siRNAs), new classes of small ncRNAs (sncRNAs) were discovered as fragments derived from canonical ncRNAs like tRNAs, small nucleolar RNAs (snoRNAs) and snRNAs (Ender et al. 2008; Lee et al. 2009; Taft et al. 2009; Chen and Heard 2013). Initially, the sncRNAs derived from other canonical ncRNAs were considered the products of random degradation (Shi et al. 2022). However, multiple lines of experimental evidence have demonstrated distinct roles in cell biology of these derived fragments involved in cancer, neurological diseases, viral infection, and others (Sun et al. 2018; Rosace et al. 2020; Coley et al. 2022; Winek and Soreq 2025).

The vault RNAs (vtRNA) are a less studied family of medium-sized ncRNAs transcribed by RNA polymerase III, which were initially described in association with the vault particle, a ribonucleoprotein particle that is ubiquitous in eukaryotes but whose function remains elusive, reviewed in (Büscher et al. 2020; Gallo et al. 2022). In humans, there are four vtRNAs located in two loci on the long arm of chromosome 5: locus vtRNA1 (5q31.3) with vtRNA1-1, vtRNA1-2, and vtRNA1-3, and locus vtRNA2 (5q31.1) with vtRNA2-1. Notably, only approximately 5% of vtRNAs are associated with the vault particle, whereas the remaining ∼95% are free in the cytoplasm, and their function remains poorly understood. Some reports have suggested that vtRNAs are potential precursors of functional small-derived fragments, which could exert regulatory functions through the miRNA pathway (Persson et al. 2009; Pillai et al. 2010; Miñones-Moyano et al. 2013; Hussain et al. 2013; Kong et al. 2015; Sajini et al. 2019; Fort et al. 2020; Seal et al. 2020; Alagia et al. 2023). The nomenclature used in the literature to refer to vtRNA-derived fragments is highly variable; therefore, throughout this work we use the term “small vault RNAs” with the abbreviation “svtRNAs,” as also adopted in a recent review on the biogenesis and function of sncRNAs in animals (Jouravleva and Zamore 2025).

The first report mentioning the possibility that vtRNA-derived fragments could function as miRNAs dates back to a 2009 study from Carlos Rovira’s group (Persson et al. 2009). Specifically, they showed that vtRNA1-1 could be processed into two dominant fragments (svtRNAa and svtRNAb) in the MCF-7 cell line, which were dependent on DICER and independent of DROSHA. Additionally, they reported the possibility of fragments binding to the Argonaute proteins (AGO2 and AGO3) and that svtRNAb could directly modulate the repression of the CYP3A4 protein. This was also confirmed in the HepG2 cell line (Meng et al. 2016). More recently, HITS-CLIP (High-throughput sequencing of RNA isolated by crosslinking immunoprecipitation) assays also confirmed that vtRNAs can bind proteins involved in miRNA biogenesis and Argonaute proteins (Wang et al. 2020). Related to the generation and processing of vtRNAs into small fragments, Michaela Frye’s group demonstrated that the protein NSUN2 (NOP2/Sun RNA methyltransferase 2) was shown to be capable of binding and methylate specific bases of vtRNA1-1, vtRNA1-2, and vtRNA1-3, leading to changes in the abundances and identities of the svtRNAs produced (Hussain et al. 2013). Specifically, svtRNA4, derived from the 3′ end of vtRNA1-1, changes its abundance in response to cytosine 69 methylation, and this impacts the regulation of CACNG7 and CACNG8 (calcium voltage-gated channel auxiliary subunit gamma 7/8), genes directly regulated by svtRNA4 (Hussain et al. 2013) through the miRNA pathway. The same group also showed that NSUN2-mediated regulation of vtRNA1-1 processing to svtRNA4 is important for fibroblast differentiation (Sajini et al. 2019). More recently, Gullerova’s group demonstrated that a svtRNA derived from vtRNA1-2 associates with the AGO protein, localizes predominantly to the nucleus, and its deletion is capable of impaired proliferation and membrane-associated genes (Alagia et al. 2023).

Notably, vtRNA2-1 was initially annotated as a miRNA precursor (hsa-mir-886) in version 10 of the miRBase database (August 2007) (Landgraf et al. 2007). Later, it was reclassified and renamed as vtRNA2-1 due to its homology to the other vtRNAs at locus vtRNA1 (Stadler et al. 2009), being removed from the miRBase database. Several reports have studied the two dominant svtRNAs derived from vtRNA2-1, previously identified in the literature as hsa-miR-886-3p and hsa-miR-886-5p. hsa-miR-886-3p was reported to be dysregulated in cancer and associated with tumorigenesis in several tissues, particularly as a tumor suppressor in prostate cancer (Aakula et al. 2015; Fort et al. 2020), bladder (Nordentoft et al. 2012), breast (Tahiri et al. 2014), lung (Cao et al. 2013; Shen et al. 2018) and thyroid (Xiong et al. 2011). However, the same svtRNA has been reported as an oncogene in kidney (Yu et al. 2014), colorectal (Schou et al. 2014) and esophageal tissues (Okumura et al. 2016). hsa-miR-886-5p was described as a tumor suppressor in the lung (Gao et al. 2011) and liver (Han et al. 2012) and, conversely, as an oncogene in oral (Xiao et al. 2012), cervical (Kong et al. 2015; Li et al. 2011), lymphoma (Liu et al. 2013), breast (Zhang et al. 2014), thyroid (Dettmer et al. 2014), bladder (Khoshnevisan et al. 2015) and myeloma (Xiang et al. 2019) cancers. Some target genes were reported for these svtRNAs. In the case of hsa-miR-886-3p: PLK1, TGFB1, CDC6, CXCL12, PITX1 and FXN (Pillai et al. 2010; Xiong et al. 2011; Mahishi et al. 2012; Cao et al. 2013; Yu et al. 2014) and for hsa-miR-886-5p: BAX and p53 (Kong et al. 2015; Li et al. 2011).

Despite this accumulating evidence, no consensus annotation or systematic catalogue of svtRNAs currently exists. Instead, current knowledge relies on isolated studies that use distinct methodologies and focus on specific cell lines or experimental conditions. Consequently, definitions, genomic coordinates, and analytical criteria vary substantially across reports, limiting reproducibility and preventing meaningful comparisons among studies.

Here, we established an annotation pipeline based on the third-party software FlaiMapper (Hoogstrate et al. 2015), to determine a comprehensive landscape of svtRNA expression across multiple human cell lines. We developed a series of structural and expression-based criteria to filter and collapse the raw FlaiMapper output, generating two lists of svtRNA candidates. One list reflects a “miRNA-like” biotype hypothesis, while the second represents a proof-of-concept to test the possible existence of svtRNAs without the restrictions of miRNA-like feature annotation. To this end, we defined a broader “Total” (as in total cellular RNA) biotype with relaxed structural constraints, for which we do not propose a specific functional hypothesis. Using this approach, we analyzed svtRNAs in datatsets enriched for sncRNAs associated with Argonaute proteins and datasets from diverse normal primary and tumor derived human cell lines. We demonstrate that human cell lines appear to generate specific svtRNAs from vault RNAs across datasets. Several svtRNAs identified by our pipeline correspond to previously reported and experimentally validated molecules, whereas others represent novel svtRNA candidates that warrant further experimental validation and functional characterization. We also determined that some miRNA-like svtRNAs are highly abundant in human cell lines and in AGO-enriched datasets, comparable to canonical miRNAs.

## Material and Methods

### Data collection and preprocessing Summary of datasets

Online omics repositories, such as Sequence Read Archive (SRA) and Gene Expression Omnibus (GEO), were searched for datasets of human small RNAs associated with Argonaute proteins (AGO-IP/CLIP/PAR-CLIP). Two reference sets of total small RNA-seq data for normal and tumor human cell lines were also recovered from SRA. We included multiple datasets obtained via distinct methodological strategies to study the association between sncRNAs and AGO proteins. These include standard immunoprecipitation (IP), ultraviolet cross-linking followed by immunoprecipitation (CLIP), RNA immunoprecipitation (RIP), and photoactivatable ribonucleoside-enhanced UV cross-linking followed by immunoprecipitation (PAR-CLIP). All libraries were prepared using the Illumina TruSeq kit for subsequent sequencing on Illumina platforms.

The data retained for analysis are summarized in Table 1. See Figure 1 and Supplementary Table 1 for detailed summaries of pipeline, datasets and accession numbers. Raw FASTQ files were downloaded from the selected SRA or GEO datasets.

**Figure 1.**
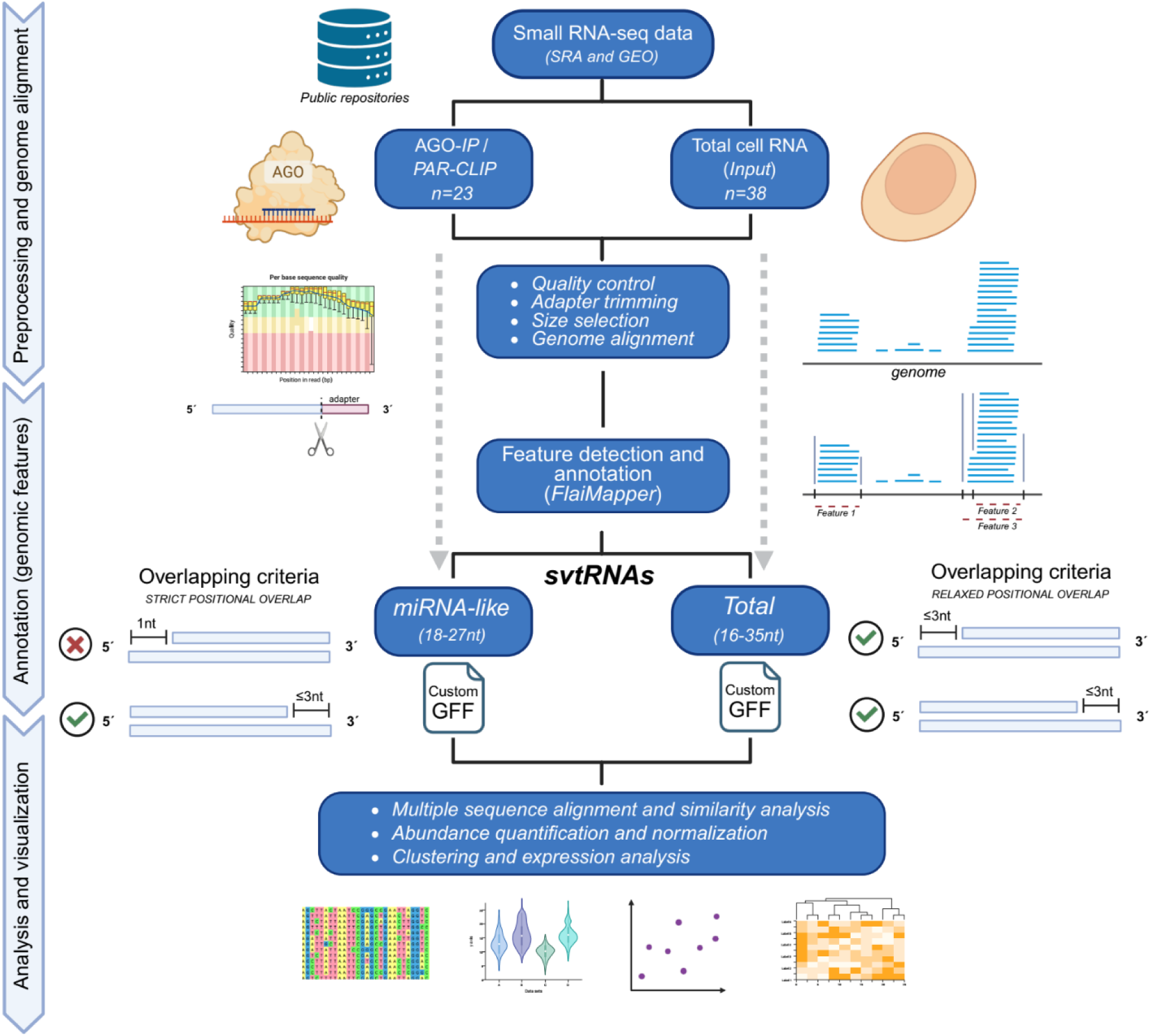
Overview of the svtRNA annotation pipeline. Schematic representation of the workflow used to identify and classify svtRNAs (created in BioRender). Two list of svtRNAs were annotated: (i) a *miRNA-like* set approach based on AGO-IP datasets and restricted to fragments of 18–27 nt with AGO association, and (ii) a *Total* set approach based on total small RNA-seq libraries of normal cell lines without constraints related to miRNA biology. Both strategies involve FlaiMapper-based prediction, filtering, structural evaluation, and collapsing of 5′ and 3′ isomiRs to generate svtRNA annotations in GFF format. The miRNA-like set represents the primary annotation proposed for svtRNA quantification, whereas the Total set serves as a proof-of-concept to explore vtRNA-derived fragments under relaxed constraints.

**Table 1.** Summary of datasets selected of small RNA-seq data for analysis.

| Dataset Type | Total Samples (n) | Cell lines (n) | Tissues (n) |
| --- | --- | --- | --- |
| *AGO IP/PAR-CLIP* | 23 | 12 | 8 |
| *Normal cell lines* | 38 | 38 | 18 |
| *Tumor cell lines* | 59 | 59 | 9 |

### Preprocessing pipeline

Raw FASTQ files were first explored using FastQC to detect contamination with adapter sequences expected after standard small RNA-seq library preparation protocols. Adapter sequences were eliminated using the software Cutadapt (v. 3.2) (Martin 2011) with the following command, establishing a minimum length of 16 nucleotides and a maximum of 100 nucleotides for retained sequences: ‘cutadapt -a [adapter sequence] -m 16 -M 100 -o <output.file> <input.file>’. Filtered reads were mapped to the current human genome assembly (hg38) using Bowtie2 (v. 2.3.5.1) (Langmead and Salzberg 2012) with default parameters.

The output SAM files were indexed with Samtools (v. 1.10) (Danecek et al. 2021) for subsequent manipulations. The statistical data related to the preprocessing of this data, including adapter trimming results and percentage of reads mapped to the human genome, are summarized in Table 2. Finally, the ordered and indexed SAM files were used as input for annotation of svtRNAs as described in the following sections (Figure 1).

**Table 2.**
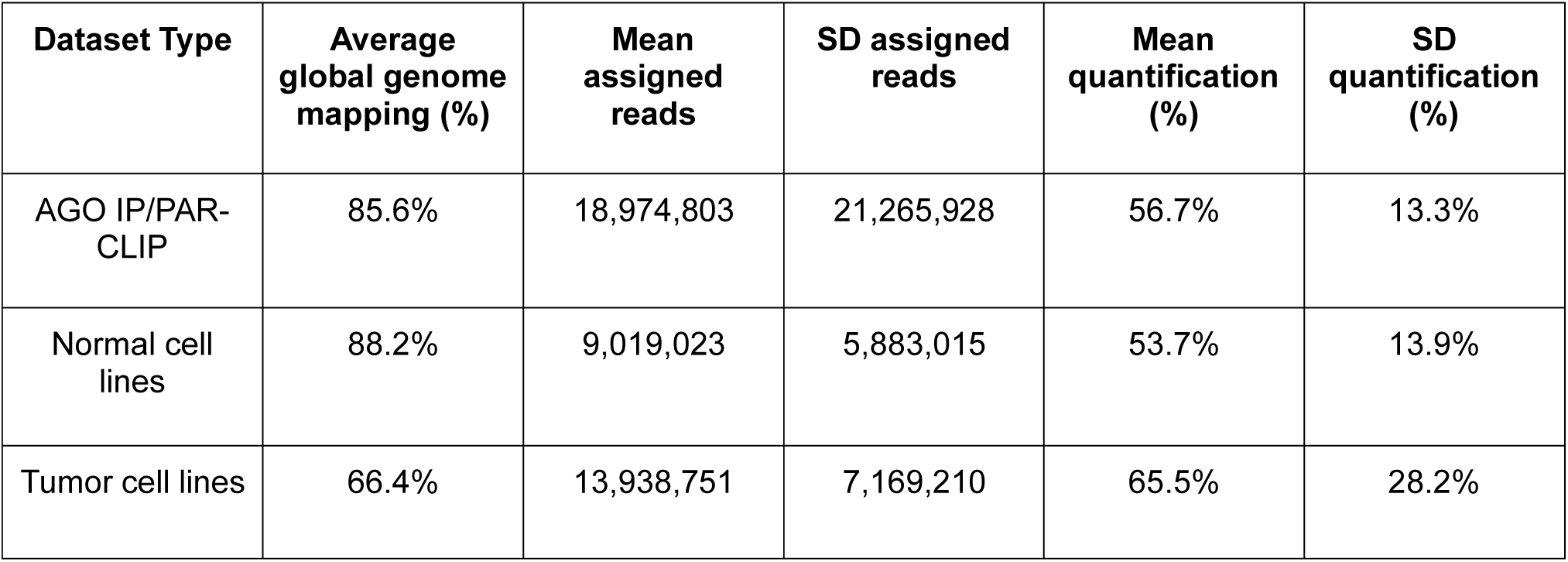
Summaries of the statistics of mapping to genome of selected small RNA-seq datasets.

### Annotation pipeline for search, filtering, and refinement of small RNAs derived from vtRNAs. Detection of sncRNA fragments using FlaiMapper

The next processing step was the annotation of svtRNAs using the “Fragment Location Annotation Identification Mapper” (FlaiMapper) software (Hoogstrate et al. 2015). Briefly, FlaiMapper defines a fragment as the region in a precursor ncRNA between the most frequent start and end positions, empirically determined from reads mapped to the precursor ncRNA’s genomic coordinates. After reconstruction of the final detected sncRNA fragments, the output of FlaiMapper is a list of detected fragments in General Feature Format (GFF3), consisting in an identifier automatically generated during the annotation step, genomic coordinates for the start and end position of each fragment, and a score calculated as: the density of reads supporting a given fragment, penalized by distance to the fragment’s expected start and end positions, as calculated from the total number of reads mapped to those coordinates in the genome.

### Annotation of small vault RNAs and isoforms

The raw output of FlaiMapper was further processed using *in-house* Bash and Python scripts to feed a second annotation step, retaining fragments derived from genomic regions corresponding to vtRNAs. Pseudogene-derived vtRNAs were excluded from the final analysis. The maximum length of 35nt for the group of Total svtRNAs reflects the insert size expected for the human normal primary cell line dataset according to the library preparation protocol (McCall et al. 2017). Identical fragments detected redundantly across samples were collapsed by summing their FlaiMapper scores. After selecting fragments whose coordinates lay entirely within a ±2-nt margin of the annotated vtRNA genomic interval, detected svtRNA fragments were ordered by genomic coordinates and scanned sequentially. To construct the miRNA-like svtRNA set from small RNA sequencing data from AGO association assays, overlapping candidates were defined as belonging to the same 5′ group when they shared an identical 5′ coordinate, and differed by at most 3 nt at their 3′ end, retaining the longest fragment to represent each 5′ cluster; subsequently, fragments with lengths outside the canonical miRNAs (18–27 nt) were excluded (Figure 1). Total svtRNAs set were annotated from normal human primary cell line data, allowing lengths between 16 and 35 nt, and redefining the conditions of overlapping fragments to allow adjacency by up to 3 nt at both 5′ and 3′ ends. After collapsing svtRNA candidates, fragments with an aggregated FlaiMapper score ≥100 were retained and merged with the most recent miRBase annotation (v. 22) (Kozomara et al. 2019), and exported as two GFF3 files to enable separate quantification of miRNA-like and Total svtRNA sets candidates (https://github.com/functional-genomics-section/svtRNA-annotation/blob/main/svtRNA_annotation_v1.gff3).

### Feature quantification

Human miRNAs and annotated svtRNA candidates (miRNA-like and Total sets) were quantified from aligned reads using *featureCounts* (Subread v2.0.1) software. Quantification was performed considering partially overlapping fragments of near-ambiguous or ambiguous origin (see Subread documentation). For each dataset, *featureCounts* produced a raw read count matrix for all annotated miRNAs and svtRNAs. The parameters used were: ‘ --minOverlap 16 --fraction -O --fracOverlap 0.9 --primary -M -s 1’.

### Normalization and feature abundances

Raw counts were normalized to reads per million (RPM) to account for differences in sequencing depth across libraries. For each sample, RPM values were computed by dividing each feature’s count by the total number of mapped reads assigned to annotated small RNAs in that sample and multiplying by 10^6^. To obtain a single representative abundance value per feature within each experimental condition (e.g., AGO association, normal cell lines, tumor cell lines), RPM values were averaged across samples (for each dataset) and across datasets (for all conditions) to yield dataset-level and condition-level mean RPM per feature. This mean RPM value served as the primary quantity used for compositional summaries (percentages) and for comparisons between svtRNA and miRNA distributions, as well as to analyze svtRNA abundance according to vtRNA of origin, positional encoding (5’ end, 3’ end, internal), etc. Mean RPM values were transformed using log10(RPM+1) for visualization purposes (Supplementary Table 2).

To determine the relative contribution of each svtRNA to the total svtRNA signal as defined by the reads associated with annotated svtRNAs, svtRNA abundances were extracted for each condition, and each fragment’s contribution to the total svtRNA signal in that condition was computed as the mean RPM of that feature, divided by the sum of mean RPM values for all svtRNA features in that condition. For within-family composition (precursor *family*: svtRNAs derived from one vtRNA), the same calculation was applied within each vtRNA family grouping, i.e., dividing each fragment’s mean RPM by the sum of mean RPM values for svtRNAs derived from that family. For pooled summaries across normal and tumor cell lines, condition-level mean RPM values were calculated and averaged for each feature, prior to computing composition-based percentages as above (Supplementary Table 3).

### Statistical analysis and data visualization

All statistical analysis and downstream data visualizations were performed using the R statistical computing language (v. 4.5.1) in an RStudio environment (v. 2025.09.2). The *dplyr, readr,* and *tidyverse* packages were used for manipulation of read count matrices and to calculate basic descriptive statistics (quartiles, mean, median, frequency distribution). Separate expression matrices were constructed according to each condition (AGO-association datasets, normal cell lines, tumor cell lines). svtRNA features were identified in count matrices by pattern matching of the feature identifier and assigned to one of four “families” according to vtRNA precursor (svtRNA1-1, svtRNA1-2, svtRNA1-3, svtRNA2-1). miRNA and svtRNA features with zero expression across all samples within a condition were removed.

### Global abundance comparisons of svtRNAs versus miRNAs

To compare svtRNA abundance to the background miRNA population within each dataset/condition, feature-level abundance values were summarized across samples and visualized as distribution plots. For these global distribution comparisons, RPM values were transformed using log10(RPM + 1) to stabilize variance and accommodate the dynamic range typical of small RNA-seq data. For each condition, the resulting feature-level log-transformed values were visualized using violin plots to represent the overall distribution of expressed sncRNAs (miRNAs and svtRNAs), with svtRNA points overlaid using Sina/jittered point representations and colored according to vtRNA precursor family. Mean and median miRNA expression values were overlaid as reference lines to provide descriptive distributional context in each dataset.

### Compositional summaries of svtRNA signal and treemap visualizations

To visualize the relative contribution of individual svtRNA fragments within each condition, compositional summaries were computed from mean RPM values. Briefly, svtRNAs were extracted, and each fragment’s percentage contribution to the total svtRNA signal was computed as described in methods. These fragment-wise percentages were visualized using treemap plots (treemapify library), with rectangle area proportional to mean RPM and fill color indicating vtRNA precursor family.

### Heatmaps and svtRNA clustering

svtRNA expression patterns across small RNA-seq samples from normal and tumor cell lines were visualized using heatmaps with the *heatmap.2* package. Heatmap matrices were built from log-transformed abundance values. Hierarchical clustering was performed on both rows (svtRNAs) and columns (samples) using a Euclidean distance metric and default linkage criteria to generate dendrograms cluster analysis of svtRNA features according to their abundance across samples, and of small RNA-seq samples according to the distribution of svtRNA abundances within each sample. To focus on consistently expressed and relatively abundant fragments, a quartile-based filtering strategy was applied, retaining only svtRNAs in the upper quartile of expression for at least 10% of small RNA-sequencing samples for that condition (normal cell lines = 4, tumor cell lines = 6). Row-side color annotations were used to indicate the vtRNA precursor family for each svtRNA.

### Intersection and length-abundance analysis

For the miRNA-like svtRNA set analyzed across multiple independent AGO-association datasets, an UpSet plot was generated to summarize the overlap of svtRNAs exceeding a defined abundance threshold across datasets. A svtRNA was considered “abundant” if its expression was above the average miRNA abundance for that dataset. The percentage of AGO-associated svtRNA signal corresponding to each fragment was displayed as a bar chart above the intersection matrix, and an additional bar chart summarized the number of abundant svtRNAs detected per dataset.

To relate svtRNA abundance to fragment length and positional origin within vtRNA precursors, fragment lengths (nt) were computed from the annotated precursor sequences. SvtRNAs were assigned to one of three positional categories based on precursor-relative location: 5′-end, 3′-end, or internal (defined as fragments within 25 nt of the corresponding precursor end for 5′/3′ categories, and internal otherwise). Using the datasets from which each group of svtRNAs were annotated (AGO-associated datasets for the miRNA-like group and normal cell line data for the Total group), each fragment’s percentage contribution to total svtRNA signal was calculated, and these values were displayed as scatter plots of fragment length versus percent svtRNA abundance, with point color encoding vtRNA family and point shape encoding positional category. In addition, the total svtRNA signal attributable to each positional category was summarized in a bar plot. Pie charts were generated to display the proportion of detected svtRNA fragments belonging to each positional category within each vtRNA precursor family.

### Multiple sequence alignments and inspection of coverage

Multiple sequence alignments were generated using vtRNA precursor sequences together with their corresponding annotated svtRNA fragments (miRNA-like and Total sets), both before and after collapsing of overlapping isoforms (Supplementary Figure 1). Alignments were constructed using Jalview (v. 2.11) with the Clustal Omega alignment algorithm (default parameters) (Sievers and Higgins 2021) to visualize sequence conservation, fragment boundaries, and positional relationships between svtRNAs and their precursor vtRNAs. In parallel, read coverage and alignment patterns at vtRNA genomic loci were manually inspected using the Integrative Genomics Viewer (IGV) (v. 2.16) (Thorvaldsdóttir et al. 2013), with the human genome assembly hg38 as the reference. Per-nucleotide coverage across vtRNA loci was calculated from previously generated BAM files corresponding to normal primary and tumor human small RNA-seq datasets using SAMtools (samtools depth) (Danecek et al. 2021), restricting the analysis to annotated vtRNA genomic coordinates. Coverage values were normalized within each vtRNA in each sample by expressing per-base coverage as a percentage of the total coverage across the full-length vtRNA. Coverage profiles per vtRNA were visualized in R using ggplot2.

## Results

### Small vault RNA-derived fragments (svtRNAs) are generated across human cell lines

To characterize and classify fragments derived from human vtRNAs, we implemented two strategies. Several studies have suggested that some svtRNAs may act as miRNA-like regulatory molecules, therefore we reasoned that AGO-enriched human datasets could reveal a reproducibly processed fraction of vtRNA-derived fragments. We defined a “miRNA-like” set with fragments with minimal features consistent with canonical miRNA biology, Argonaute association, and size restrictions (18-27nt) (Bartel 2018). The miRNA-like set represents the primary annotation proposed for svtRNA quantification, prioritizing fragments compatible with canonical miRNA biology. In contrast, the Total set was generated as a proof-of-concept to explore whether additional vtRNA-derived fragments that may not be associated with Argonaute proteins could be detected under relaxed constraints, size selection (18-35nt) (Figure 1 summarizes overall workflow and see methods section).

For the miRNA-like set, we analyzed 23 small RNA-seq data samples from Argonaute immunoprecipitation (AGO-IP) assays derived from 12 different human cell lines (Supplementary Table 1). The annotation strategy led us to define an initial set of svtRNA candidates, where we applied a collapsing step for posterior quantification and robust study of miRNA-like svtRNAs. This step accounts for natural 5′ (±1 nt) and 3′ (±3 nt) variation defined for isomiRs (Hoogstrate et al. 2015; Bofill-De Ros et al. 2020). After this collapsing step, we obtained 17 miRNA-like svtRNAs (see alignment in Supplementary Figure 1), a final annotation resource of miRNA-like svtRNAs constituting a GFF3 file (available at https://github.com/functional-genomics-section/svtRNA-annotation/blob/main/svtRNA_annotation_v1.gff3) used for the following quantification analysis.

### Abundance patterns of miRNA-like svtRNAs in AGO-IP datasets

Across the AGO-IP dataset, we observed that, on average, 3 svtRNAs per set had abundances higher than the average miRNA abundance. This number varied across datasets: GSE43666 Cervix (HeLa) exhibited 9 abundant svtRNAs, whereas others, such as GSE56836 Kidney (HEK293), have none (Figure 2A). These differences likely reflect variation in vtRNA (precursor) expression, processing efficiency, stability, and/or AGO loading preferences among cell lines, as well as potential technical differences between datasets. Notably, among the three cervix-derived datasets (HeLa cell lines), two of them (GSE29116 and GSE45506) displayed only two svtRNAs, both derived from vtRNA2-1, above the average miRNA distribution, whereas the third dataset (GSE43666) showed nine svtRNAs originating from all vtRNAs (Figure 2A). This difference could also anticipate variations in svtRNA abundance linked to the specific Argonaute proteins (AGO1-4) captured in each assay, as was shown for miRNAs (Dueck et al. 2012). Specifically, in the GSE43666 dataset, only AGO2 was immunoprecipitated in the assay, while in GSE29116 and GSE45506, all four Argonautes (AGO1–4) were pulled down.

**Figure 2.**
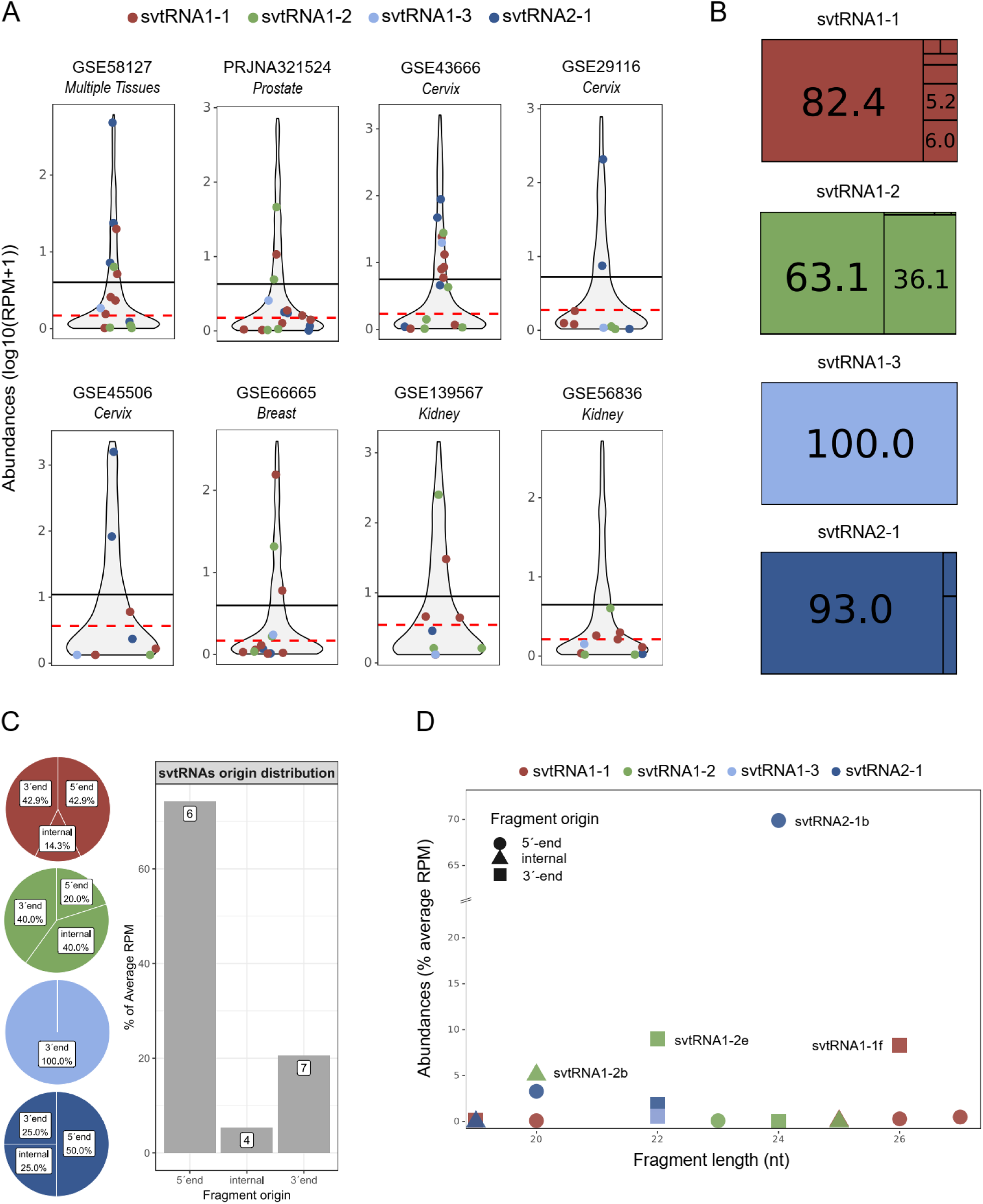
Abundance and characteristics of the miRNA-like svtRNAs identified in AGO-IP datasets. **A.** Sina plots showing the distribution of 17 svtRNA abundances across AGO-IP datasets, log10-transformed read counts are shown (5-95th percentile). Each dot represents a svtRNA in one dataset; colors correspond to the vtRNA precursor. The black line indicates the global mean miRNA abundance, and the red line marks the median for each dataset. **B.** Treemap summarizing the relative abundance of svtRNAs within each vtRNA precursor (percentage number of svtRNAs with less than 5% of reads are not shown). **C.** Pie chart of the distribution of svtRNAs according to their positional origin within vtRNAs (5′ end, internal, or 3′ end), shown as different fragments (left) and bar plot of abundance proportions (right). **D.** Dot plot integrating svtRNA abundance, fragment length, and positional origin in AGO-IP datasets.

The number of miRNA-like svtRNAs annotated per vtRNA precursor is variable: vtRNA1-1 yielded seven svtRNAs, vtRNA1-2 yielded five svtRNAs, vtRNA1-3 yielded one svtRNA and vtRNA2-1 yielded four svtRNAs (Figure 2B). The abundance of the svtRNAs annotated for each vtRNA precursor is heterogeneous. Each vtRNA gene precursor gives rise to a dominant fragment that accounts for more than half of the total svtRNAs derived from it (63.1-100%, Figure 2B). The average svtRNA length of the miRNA-like svtRNAs is 22.7 ± 2.6 nt, comparable with the average value of canonical human miRNAs (Bartel 2018) (Supplementary Table 3). Regarding positional origin within the precursor, svtRNAs are derived from: 5′-end, internal and 3′-end regions of vtRNA, showing that svtRNAs are processed without a clear pattern of positional origin (Figure 2C). If we consider all of the vtRNA precursors, there are slightly more svtRNAs from 5′ (6) and 3′-end (7) relative to internal (4) positions, with 5′-end (74.2%) being markedly more abundant than 3′-end (20.5%) and internal (5.3%) svtRNAs (Figure 2C). Combined analysis of length, origin and abundances of the svtRNAs revealed no specific enrichment on particular lengths or size for svtRNAs (Figure 2D). It is important to note that svtRNA2-1b is globally the most abundant svtRNA, accounting for 69.9% of total svtRNAs abundance. Also, the most abundant svtRNAs for each vtRNA were svtRNA1-1f (26 nt, 3′-end), svtRNA1-2e (22 nt, 3′-end), svtRNA1-3a (22 nt, 3′-end), and svtRNA2-1b (24 nt, 5′-end) (Figure 2D and Supplementary Table 3).

### Identification of highly abundant miRNA-like svtRNAs

To focus on highly abundant miRNA-like svtRNAs that could be relevant to cellular function, we selected the svtRNAs that: (i) surpassed the average for miRNA abundances in at least one AGO-IP dataset and (ii) were in the upper quartile of abundance in at least 10% of primary and tumor cell line samples (Supplementary Figure 2). This ensures that the selected svtRNAs are abundant in Argonaute proteins in at least one cell line and are also abundant in total human cell lines. Five miRNA-like svtRNAs satisfied these criteria (Figure 3A): one from vtRNA1-1 (svtRNA1-1f), one from vtRNA1-2 (svtRNA1-2e), and three from vtRNA2-1 (svtRNA2-1a, svtRNA2-1b, and svtRNA2-1d) (Figure 3A). Remarkably, four of the five highly abundant miRNA-like svtRNAs were independently reported by experimental studies and were validated by Northern blot and/or RT-qPCR by different groups (Persson et al. 2009; Pillai et al. 2010; Treppendahl et al. 2012; Miñones-Moyano et al. 2013; Hussain et al. 2013; Kong et al. 2015; Sajini et al. 2019; Alagia et al. 2023) (Supplementary Table 3). Of note, another svtRNA derived from vtRNA1-1 (svtRNA1-1e) is highly abundant in normal and tumor cell line total small RNA-seq but does not surpass the average for miRNA abundances in the analyzed AGO datasets (Figure 3A), supporting the idea that not all cell-abundant svtRNAs are also associated with AGO compartment. In contrast, svtRNA1-1g and svtRNA1-1b are abundant in two AGO datasets but show low abundance in both normal and tumor cell line total small RNA-seq (Figure 3A). This may be related to differences in the enrichment preferences of svtRNA loading to the different AGO proteins.

**Figure 3.**
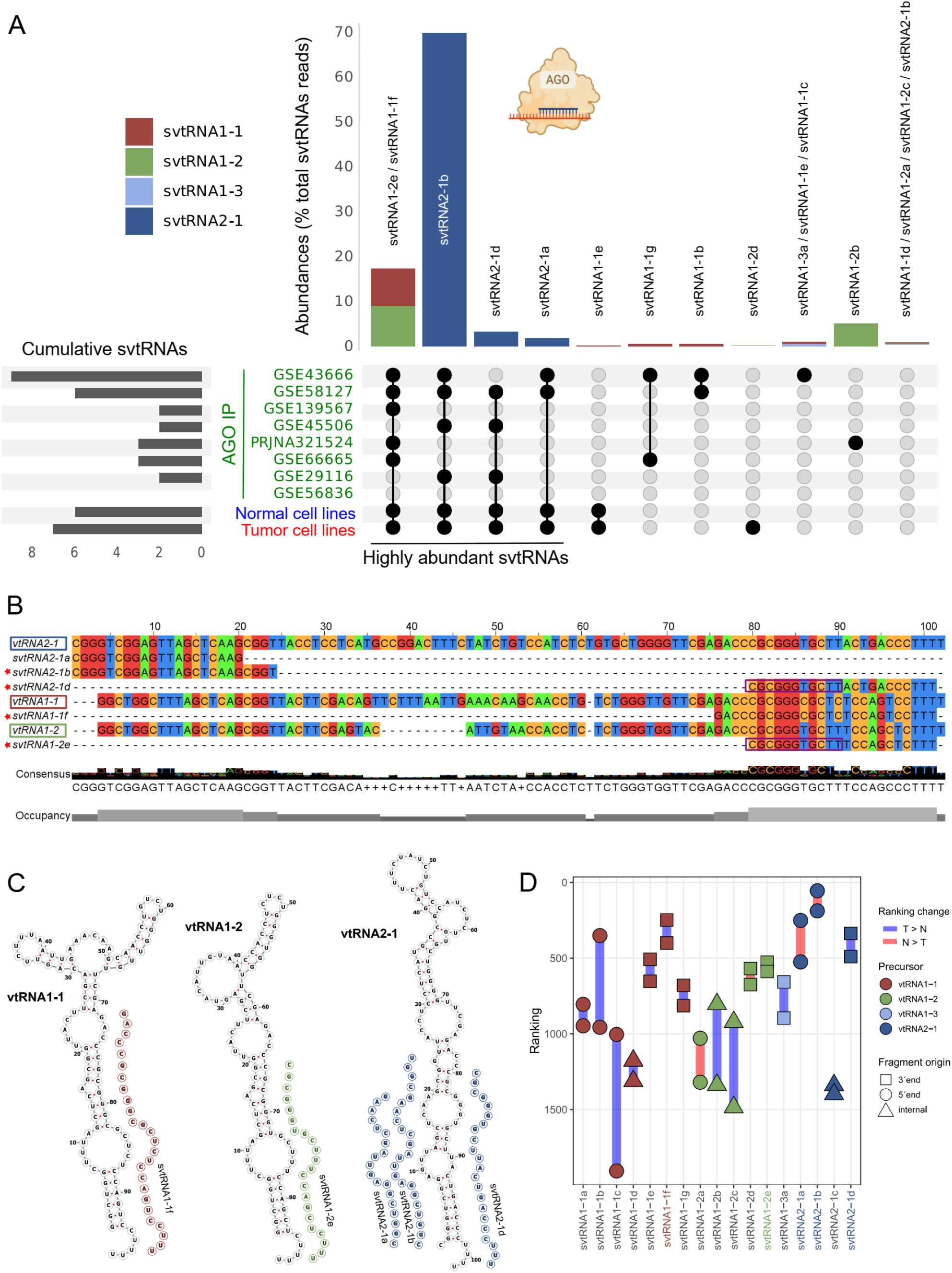
Highly abundant miRNA-like svtRNAs across AGO-IP datasets. **A.** Upset plot summarizing abundance criteria used to define five highly expressed miRNA-like svtRNAs. Bars indicate the number of datasets in which each svtRNA exceeded the mean miRNA abundance (AGO-IP) or the first quartile of miRNA abundance (Normal and Tumor cell lines). The svtRNAs that surpass the average miRNA abundance in at least one AGO-IP and the first quartile of 10% of primary and/or tumor cell lines are selected as highly expressed svtRNAs. **B.** Sequence alignment of highly abundant svtRNAs. Red stars indicate fragments previously validated experimentally by Northern blot or RT-qPCR. Shared nucleotides, including seed-like regions between svtRNA2-1d and svtRNA1-2e, are boxed in purple. **C.** The predicted secondary structure of the precursor vtRNAs and the location of the highly svtRNAs are shown. **D.** Ranking changes of miRNA-like svtRNAs when comparing normal vs tumor cell lines. Each dot represents a svtRNA; colors indicate vtRNA precursor; shapes indicate positional origin. Blue lines represent increases in rank between normal to tumor datasets; red lines indicate decreases in rank.

A more detailed examination of svtRNAs, sequence alignments, and predicted structures highlighted biological features of interest (Figure 3B–C). svtRNA2-1b and svtRNA2-1d have partial complementary and resemble the typical miRNA duplex (Figure 3C). Another aspect to remark upon is that svtRNA2-1d and svtRNA1-2e share the first 11 nucleotides, including the seed region, suggesting the possibility of partially overlapping regulatory targets, an aspect not previously reported or investigated in prior studies analyzing individual svtRNAs (see purple boxes in Figure 3B).

The previously reported roles for svtRNAs in cancer (Büscher et al. 2020; Yeganeh and Hernandez 2020; Gallo et al. 2022) led us to explore changes in their abundances between normal and tumor cell lines by examining miRNA and svtRNA rank changes (Figure 3D). We observed that there is a global increase in the ranking (blue lines) of svtRNAs between normal and tumor cell lines (13/17 svtRNAs), which may reflect differences in the relative abundances of svtRNAs between these conditions. Only four svtRNAs (derived from vtRNA1-2 and vtRNA2-1) decreased in rank in tumor cell lines compared to normal cell lines (red lines, Figure 3D). Remarkably, the svtRNA derived from the 5′-end of vtRNA2-1 (miRNA-like svtRNA2-1b) is ranked comparably to the most abundant miRNAs, achieving the 55^th^ position in tumor samples, which is exceptionally high (Figure 3D).

### Total svtRNAs set could be defined from total small RNA-seq datasets

As a proof of concept, and to avoid overlooking abundant svtRNAs that may not be associated with Argonaute proteins, we generated a svtRNA annotation using relaxed miRNA-related criteria and total cellular small RNA-seq datasets, hereafter referred to as the Total svtRNA set. This approach enabled us to assess how svtRNA repertoires differ when annotated from small RNA-seq data using distinct analytical strategies and stringency levels. This data-driven *de novo* svtRNA set was based on a representative total small RNA-seq dataset of normal human cell lines (PRJNA358331: 38 samples from 38 different cell lines, Supplementary Table 1). After annotation (see details in methods and Figure 1) we identified 13 Total svtRNAs: four from vtRNA1-1, five from vtRNA1-2, and four from vtRNA2-1 (Supplementary Figure 1; Supplementary Table 1). No svtRNAs were identified from vtRNA1-3 in this dataset. In the Total set (13) less svtRNAs were defined than the miRNA-like set (17), probably due to relaxing the constraints for collapsing isoforms with regards to fragments that presented overlapping 5’ and 3’ ends. Nine of the 13 svtRNAs surpass the average miRNA abundances in normal cells, showing that svtRNAs potentially constitute a non-negligible component of the small RNA landscape in non-transformed cell lines (Supplementary Figure 3A). One dominant svtRNA from vtRNA1-1 (svtRNA1-1d with 67.6%) and vtRNA2-1 (svtRNA2-1a with 84.3%) is observed, whereas vtRNA1-2 presents two almost equally abundant svtRNAs (svtRNA1-2e with 38.6% and svtRNA1-2a with 38.3%) (Supplementary Figure 3B).

Complementarily, we analyzed the abundances of the Total svtRNA set in tumor cell lines (SRP109305: 59 samples from 59 distinct cell lines; Supplementary Table 1 and 2) and observed that only five svtRNAs surpass the average miRNA distribution (Supplementary Figure 3C). In the normal human cell lines svtRNA abundances within precursors showed dominant fragments, although less pronounced than in the tumor cell lines (Supplementary Figure 3B and 3D). A decrease in svtRNA abundance relative to miRNA abundances in comparison to normal cells lines may reflect a lower expression, differential processing or higher instability of the svtRNAs in the tumor cell lines (Supplementary Figure 3A and 3C). Additionally, tumor cell lines showed a stronger dominance of single svtRNAs per precursor compared to normal cell lines (Supplementary Figure 2D); vtRNA1-1 (svtRNA1-1d with 68%), vtRNA1-2 (svtRNA1-2e with 89.6%) and vtRNA2-1 (svtRNA2-1a with 99.7%)). This is similar to what was observed for the miRNA-like svtRNAs set in the AGO dataset (Figure 2B), which are also tumor cell lines. This again is consistent with the idea of differences in expression, processing or stability of the svtRNAs in tumor cell lines compared to normal cell lines (Figures 2B, 2C, 3D, Supplementary Figure 3A-D).

The average length distribution of the Total svtRNAs set is 25.9 ± 5.2 nt, longer than the average miRNA-like set (Supplementary Table 3). This could be related to the relaxed overlapping criteria. Regarding fragment origin within vtRNA precursors (5’-end, internal or 3’-end), again supports the notion that there is no preference across the different vtRNAs (Supplementary Figure 3E-F and Supplementary Table 3). Although there are slightly more svtRNAs originating from internal regions (6) compared to the 5’-end (3) and 3’-end (4), their relative abundances in normal cell lines show the opposite trend: 3’-end svtRNAs account for 43.6% of total svtRNA-associated reads, followed by 5’-end svtRNAs with 32.9%, and internal fragments with 23.4% (Supplementary Figure 3F). Remarkably, in tumor cell lines the 5’-end svtRNAs exceed the 3’-end, but the internal svtRNA population are significantly less abundant (Supplementary Figure 3F). Combined length, positional origin and abundances of the svtRNAs in normal cell lines no simple relationship can be appreciated (Supplementary Figure 3G). However, the most abundant svtRNAs appeared to be rounding the range of 22nt to 26nt (Supplementary Figure 3G). The most abundant svtRNAs for each vtRNA are svtRNA1-1d 3’-end of 26nt, svtRNA1-2e 3’-end of 26nt and svtRNA2-1a 5’-end of 24nt (Supplementary Figure 3G and Supplementary Table 3).

### Highly abundant Total svtRNAs

We identified five highly abundant svtRNAs in the Total set shared across normal and tumor datasets (Figure 4A), defined as svtRNAs that at least surpass the 1st quartile in at least 10% of total samples of normal and tumor cell lines dataset’s (Supplementary Figure 4). Among then, two derive from vtRNA1-1 (svtRNA1-1c and svtRNA1-1d), one from vtRNA1-2 (svtRNA1-2e), and two from vtRNA2-1 (svtRNA2-1a and svtRNA2-1d) (Figure 4A). Notably, four of these were identical to svtRNAs identified in the miRNA-like set: svtRNA2-1d in miRNA-like set is svtRNA2-1d in Total set, svtRNA2-1b in miRNA-like set is svtRNA2-1a in Total set, svtRNA1-1e in miRNA-like set is svtRNA1-1c in Total set and svtRNA1-1f in miRNA-like set is svtRNA1-1d Total set (Figures 3A, Figure 4 and Table 3). Also, svtRNA1-2e in the miRNA-like set is contained within svtRNA1-2e from the Total set (4 extra nucleotides at the 5’ end). The strong concordance in the precise annotation of highly abundant svtRNAs across Argonaute-associated and total cellular datasets, defined using two distinct annotation strategies (miRNA-like and Total sets), is compatible with the idea that svtRNAs are generated in a reproducible and precise manner in diverse biological contexts, tissues, etc. This agreement was observed despite differences in both the analytical stringency and the source datasets used.

**Figure 4.**
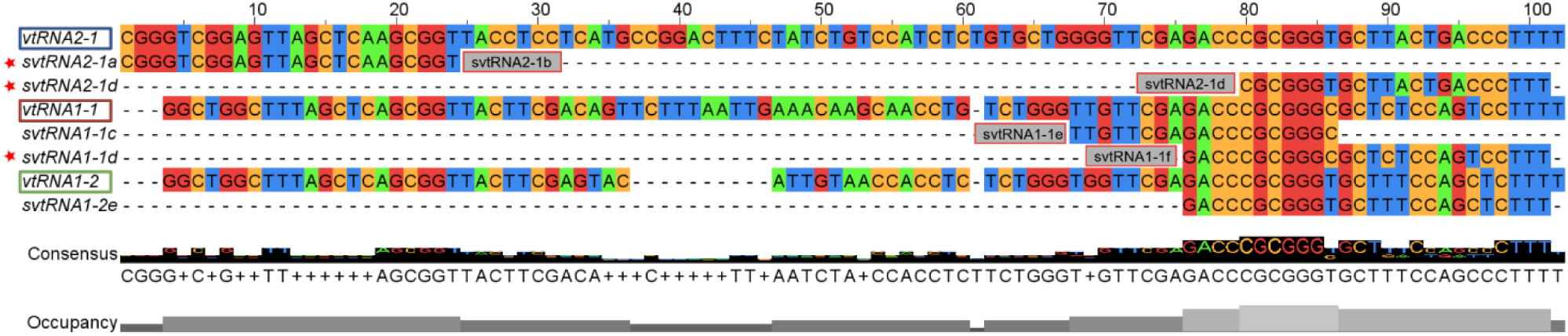
Highly abundant Total svtRNAs in normal and tumor cell lines. Sequence alignment of the highly abundant global svtRNAs. Grey boxes indicate the ID of the same svtRNA identified in the miRNA-like annotation.

**Table 3:**
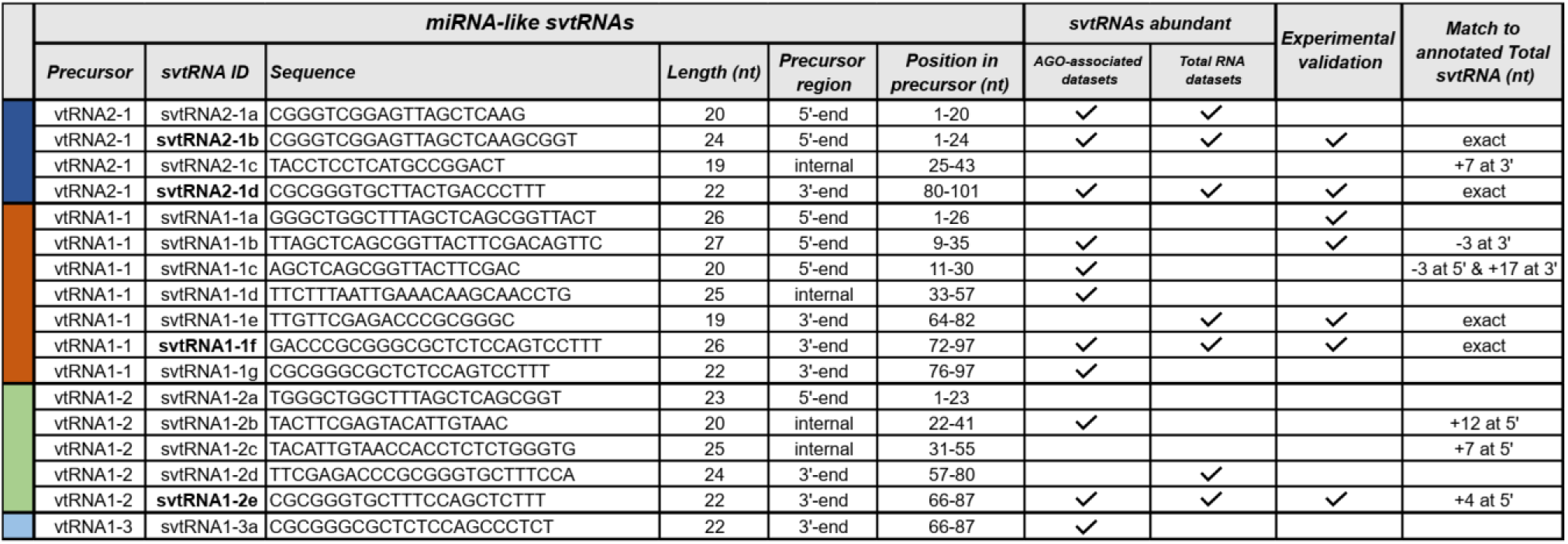
Summary of annotated miRNA-like svtRNAs. svtRNAs classified as miRNA-like are shown with their precursor, sequence, length, and position within the vtRNA. Precursor region indicates whether the svtRNA originates from the 5’-end, an internal region, or the 3’-end of the precursor. The columns under svtRNAs abundant indicate the svtRNAs that surpass the average miRNA abundance in at least one AGO-associated dataset and the first quartile of 10% of primary and/or tumor cell lines dataset. Experimental validation indicates svtRNAs previously validated experimentally by Northern blot or RT-qPCR. Match to annotated Total svtRNA indicates the relationship with sequences in the Total svtRNA annotation set, where “exact” denotes identical sequences and values such as “+n” or “−n” indicate nucleotide extensions or truncations at the indicated ends. svtRNAs IDs shown in **bold** correspond to the four svtRNAs most abundant and with experimental validation.

## Discussion

Previous studies have reported the existence, abundance, and function of specific svtRNAs in human cells, but these analyses were largely restricted to individual cell lines or experimental systems (Persson et al. 2009; Pillai et al. 2010; Treppendahl et al. 2012; Miñones-Moyano et al. 2013; Hussain et al. 2013; Kong et al. 2015; Sajini et al. 2019; Alagia et al. 2023). Here, we extend these focused observations by providing a comprehensive landscape of svtRNA expression across multiple human cell lines. These results demonstrate that human cell lines derived from different tissue origins consistently generate specific svtRNAs from vault RNAs, in a manner that is more extensive and pervasive than previously appreciated. We show that several svtRNAs are highly abundant in human cell lines. To this end, we generated an annotation resource of human svtRNAs, provided in GFF3 format to facilitate downstream analyses. It reflects a “miRNA-like” biotype hypothesis, although other functional mechanisms cannot be discarded, and it may be of interest for biomarker discovery or in other biological contexts.

Our expression-based annotation strategy successfully identified seven miRNA-like svtRNAs that were previously described in specific cell lines, providing validation for the overall approach (Table 3 and Suplementary Table 3). Notably, four svtRNAs are highly abundant miRNA-like svtRNAs (svtRNA1-1f, svtRNA1-2e, svtRNA2-1b, and svtRNA2-1d, see Table 3 and coverage plots: Supplementary Figure 5) correspond to svtRNAs reported and validated in earlier studies (Persson et al. 2009; Pillai et al. 2010; Treppendahl et al. 2012; Miñones-Moyano et al. 2013; Hussain et al. 2013; Kong et al. 2015; Sajini et al. 2019; Alagia et al. 2023). In particular, svtRNA1-1f, derived from vtRNA1-1, was initially described by Persson et al. in 2009 and validated by Northern blot in the MCF-7 cell line, where it was named svRNAa* (Persson et al. 2009). This fragment was later detected with three additional nucleotides at the 5′ end in HEK293 cells using small RNA-seq (Hussain et al. 2013; Alagia et al. 2023). Similarly, svtRNA1-2e, derived from vtRNA1-2, was observed, annotated, and validated by Northern blot in HEK293 cells, differing only by a single additional nucleotide at the 3′ end (Alagia et al. 2023). The vtRNA2-1–derived svtRNA2-1b was detected and validated in HeLa and SiHa cell lines by stem-loop RT-qPCR and sequencing (Kong et al. 2015), whereas another vtRNA2-1–derived fragment, our svtRNA2-1d, was validated by Northern blot in T24, LD419, HS5 and HS27a cells (Pillai et al. 2010; Treppendahl et al. 2012). The existence of dominant svtRNA fragments for vtRNAs is similar to processing patterns observed for other ncRNAs biotypes, such as tRNA-derived fragments and snoRNA-derived RNAs (Jouravleva and Zamore 2025).

Our results are consistent with previous observations suggesting that some svtRNAs may interact with components of the canonical miRNA pathway. Several prior studies are consistent with the hypothesis that specific svtRNAs can associate with the RISC complex and regulate target gene expression (Persson et al. 2009; Pillai et al. 2010; Xiong et al. 2011; Li et al. 2011; Mahishi et al. 2012; Hussain et al. 2013; Cao et al. 2013; Yu et al. 2014; Kong et al. 2015; Meng et al. 2016; Alagia et al. 2023). However, to our knowledge, no dedicated annotation resource for svtRNAs has been available for routine small RNA-seq data analysis. As a result, the miRNA-like regulatory potential of svtRNAs has likely been underestimated in human cell biology.

Also, the observed concordance in the annotation of highly abundant svtRNAs across Argonaute-associated and total cellular datasets when comparing the two distinct annotation strategies (miRNA-like and Total sets) (Table 3) again supports the idea that specific svtRNAs are reproducibly detected in human cells. Only three internal, lowly expressed svtRNAs were defined in the Total set without a counterpart in the miRNA-like set, further supporting the idea that the miRNA-like annotation is sufficiently comprehensive to capture the relevant expressed svtRNAs in human cells. Therefore, we recommend the miRNA-like annotation set for svtRNA quantitative analyses, as it captures the most robustly and consistently generated svtRNAs while preserving the ability to evaluate potential miRNA-related biological functions of these molecules. Importantly, this annotation strategy minimizes discrepancies in 5’-end definition, which is critical for Argonaute loading and downstream functional analyses (provided as a GFF3 annotation miRNA-like set). This resource enables a systematic, non-redundant, and reproducible quantification of svtRNAs in human cell lines, and facilitates the reanalysis of existing small RNA-seq datasets to uncover the biological relevance of svtRNAs.

It is noteworthy that certain svtRNAs (e.g., svtRNA2-1b and svtRNA1-1f) present expression levels that are comparable to canonical miRNAs in human cells. At first glance, this may appear surprising, given previous reports indicating that only ∼3–5% of vtRNAs are processed into svtRNAs. However, Lee and colleagues reported that vtRNA2-1 in HeLa cells is present at approximately 85 fmol per 5 × 10⁵ cells, corresponding to ∼10⁵ copies per cell (Lee et al. 2011). Assuming that only 3% of vtRNA2-1 molecules are processed into svtRNAs, this would yield an estimated ∼3,000 copies per cell of a specific svtRNA, a value that exceeds the average copy number of miRNAs (∼2,400 copies per cell) (Chen et al. 2005). Such abundance argues against svtRNAs being rare byproducts of vtRNA turnover and suggest that they may represent functional RNA molecules.

Although vtRNA secondary structures may partially resemble miRNA precursors, and several studies have reported that vtRNAs can be processed by DICER in a DROSHA-independent manner (Persson et al. 2009; Lee et al. 2011; Miñones-Moyano et al. 2013; Fort et al. 2020; Alagia et al. 2023), the overall efficiency of vtRNA-to-svtRNA processing appears to be low. This limited processing efficiency may reflect the fact that vtRNAs are not fully optimized as canonical miRNA precursors (Starega-Roslan et al. 2011, 2015). Structural divergence from the optimized human miRNA hairpin structure is compatible with a reduced DICER processing efficiency, resulting in the production of relatively small amounts of svtRNAs relative to precursor. However, it is conceivable that, given the high cellular abundance of vtRNA precursors in human cells, even a low processing rate could result in biologically meaningful levels of svtRNAs. This could be compatible with evolutionary hypotheses of new DICER substrates (such as miRNAs) which propose their origin as inefficiently processed precursor RNAs (Berezikov 2011).

Interestingly, we observed that highly abundant svtRNA2-1d and svtRNA1-2e, which are derived from the 3’-ends of two distinct vtRNA precursors (vtRNA2-1 and vtRNA1-2, respectively), share an identical stretch of the first eleven nucleotides. The existence of shared 5’ seed-like sequences from svtRNAs derived from distinct vtRNA precursors again is compatible with a coordinated and non-random processing. Considering their relative abundance in Argonaute-associated datasets and key principles of miRNA biology, including seed sequence mediated target recognition, these two svtRNAs may share seed-like sequences adequate for consideration as a potential miRNA family (Bartel 2009). These svtRNAs do not share seed sequences with any annotated human miRNAs in miRBase (v22), suggesting that they potentially could regulate a distinct set of target genes in human cells.

Regarding the role of svtRNAs in cancer biology, a preliminary observation is that 13 out of 17 svtRNAs experience a global increase in rank between normal and tumor cell lines (Figure 3D). It is noteworthy that, for the same vtRNA precursor, some svtRNAs show rank changes in the opposite direction between normal and tumor cell lines. This could be related to independent ratios of processing or stability of svtRNAs, as previously reported for RNA polymerase III transcripts such as vtRNAs in association with DUSP11 activity (Burke et al. 2016). However, this observation should be interpreted with caution and extended to additional datasets for confirmation, in order to avoid potential batch effects that could confound this meta-analysis.

We acknowledge several limitations of the present study that should be considered in future work. The miRNA-like svtRNA set was derived primarily from Argonaute immunoprecipitation datasets that are largely restricted to tumor-derived cell lines, which may bias detection towards svtRNAs enriched in cancer contexts while overlooking those preferentially expressed in non-tumor cells. In addition, we deliberately selected Argonaute immunoprecipitation datasets from unperturbed conditions, avoiding experiments involving miRNA overexpression, gene knockouts, cellular stress, infection, or stimulation. While this strategy minimizes confounding effects, it may also exclude condition-specific svtRNAs. Moreover, heterogeneity in Argonaute immunoprecipitation strategies (e.g., targeting AGO2 versus all AGO1-4), RNA extraction protocols, library preparation methods, sequencing platforms, and data processing pipelines likely contributes to variability in svtRNA abundance estimates across studies. These factors may partly explain the differences observed among the three HeLa Argonaute datasets analyzed here. Consequently, comparisons between normal and tumor-derived cell lines should be interpreted with caution. Future efforts incorporating larger and more homogeneous datasets, as well as additional biological contexts, could be important to refine svtRNA annotation and to identify svtRNAs that may be missed in the present study. Extending these analyses to tissue datasets will be particularly relevant to assess tissue-specific svtRNA expression patterns and to evaluate whether or not current annotations generalize beyond tumor-derived cell models. Collectively, our data support the view that svtRNAs could represent a structured and reproducibly detected components of the human small RNA landscape.

## Conclusions

In summary, our study demonstrates that human vault RNAs appear to form a discrete and reproducibly detected repertoire of small RNA fragments that can be systematically defined through expression-based annotation. By integrating Argonaute-associated and total small RNA-seq datasets under two complementary annotation strategies, we provide a standardized miRNA-like svtRNA annotation resource in GFF3 format.

The strong concordance observed across independent datasets and analytical approaches supports the notion that svtRNAs are unlikely to represent random degradation products and instead appear to correspond to specific small RNAs within the human transcriptome. Several svtRNAs are expressed at levels comparable to canonical miRNAs and include miRNA-like fragments with previous experimental validation, reinforcing the hypothesis of their biological relevance.

This resource enables the systematic inclusion of svtRNAs in small RNA-seq analyses in future studies aimed to elucidate their biogenesis, molecular mechanisms, and potential roles in human physiology and disease.

## Supporting information

Main Manuscript

## Data Availability Statement

All datasets analyzed in this study are publicly available. Detailed information, including accession numbers and repository links, is provided in the main text and Supplementary Material.

## Author Contributions

Conceptualization: RSF, MAD; Methodology: RSF, JDS, PS; Software: JDS, RSF; Validation: JDS; Formal Analysis: JDS, RSF; Investigation: JDS, RSF; Resources: RSF, MAD; Data Curation: JDS, RSF; Writing – Original Draft: RSF, JDS; Writing – Review & Editing: RSF, MAD, PS, JDS; Visualization: RSF, JDS; Supervision: RSF, MAD, PS; Project Administration: RSF; Funding Acquisition: MAD, RSF, PS.

## Acknowledgments

The authors thank the members of the laboratory for valuable discussions and technical assistance. We also acknowledge institutional support from Facultad de Ciencias, Universidad de la República, Uruguay; Instituto de Investigaciones Biológicas Clemente Estable, MEC, Uruguay. The authors thank the IT team Héctor Caporicci y Joaquín Carrique for maintaining the computational infrastructure and servers used in this study.

## Additional Supplementary Material

**Supplementary Table 1:** Summary of small RNA-seq samples and mapping statistics. List of analyzed samples, including sequencing and alignment statistics.

**Supplementary Table 2:** Normalized expression values of svtRNAs and miRNAs. Normalized counts for svtRNAs and miRNAs across all analyzed samples.

**Supplementary Table 3:** Sequence and annotation information for svtRNA datasets. Sequence information, IDs, length, and associated statistics for svtRNAs included in the miRNA-like set and the Total set.

**Supplementary Figure 1.**
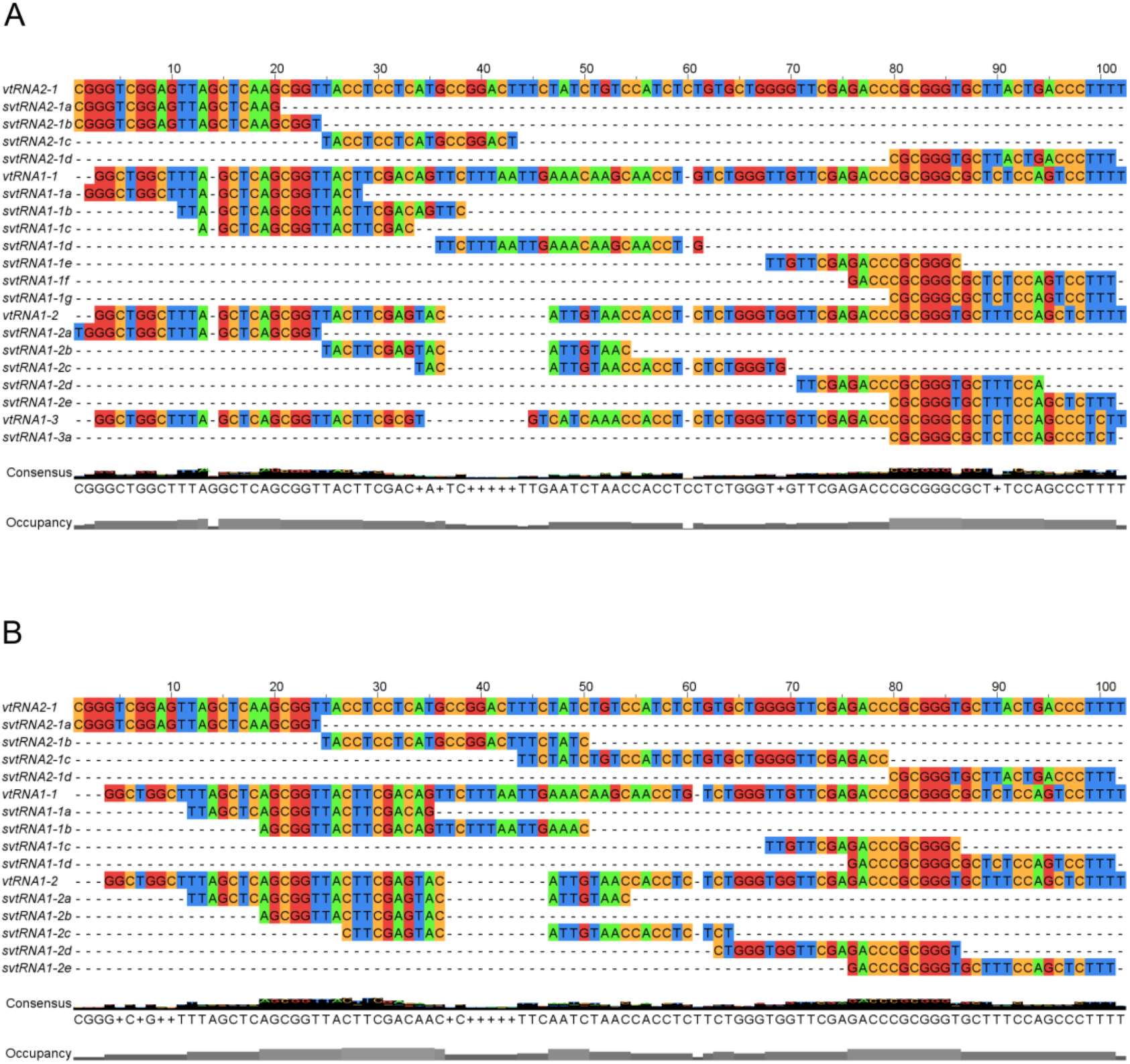
Sequence alignment of annotated svtRNAs. Multiple sequence alignment of the annotated svtRNAs identified across both strategies (**A**) miRNA-like set and (**B**) Total set. Nucleotide positions are shown relative to the vtRNA reference sequence. The consensus sequence is displayed below the alignment, along with nucleotide occupancy across positions.

**Supplementary Figure 2.**
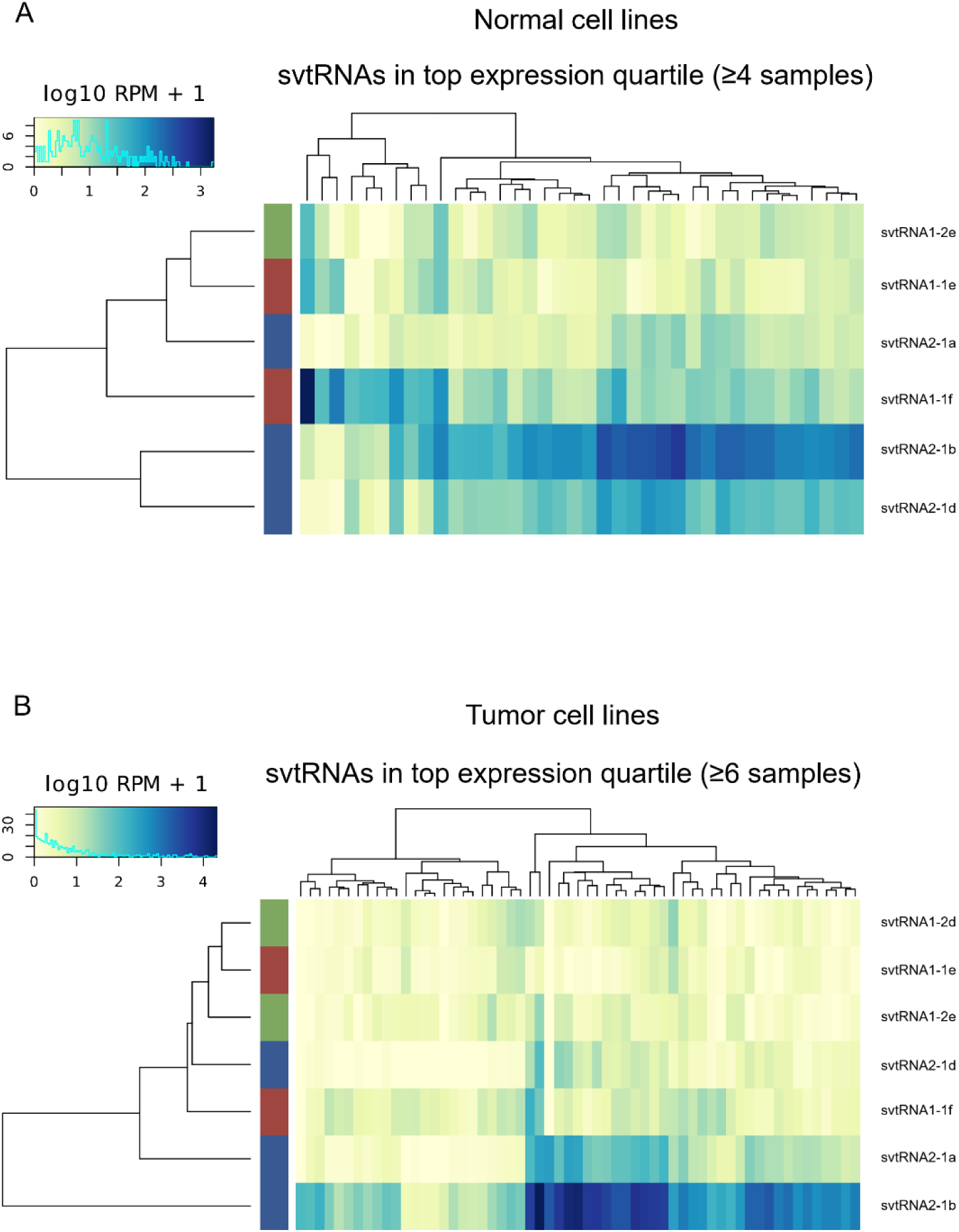
Hierarchical clustering of highly expressed miRNA-like svtRNAs in normal and tumor cell lines. Heatmaps show svtRNAs in the upper expression quartile detected in ≥4 normal samples (A) or ≥6 tumor samples (B). Values correspond to log10(RPM+1). Clustering was performed using Euclidean distance on both rows and columns. Colored side bars denote vtRNA precursor families.

**Supplementary Figure 3.**
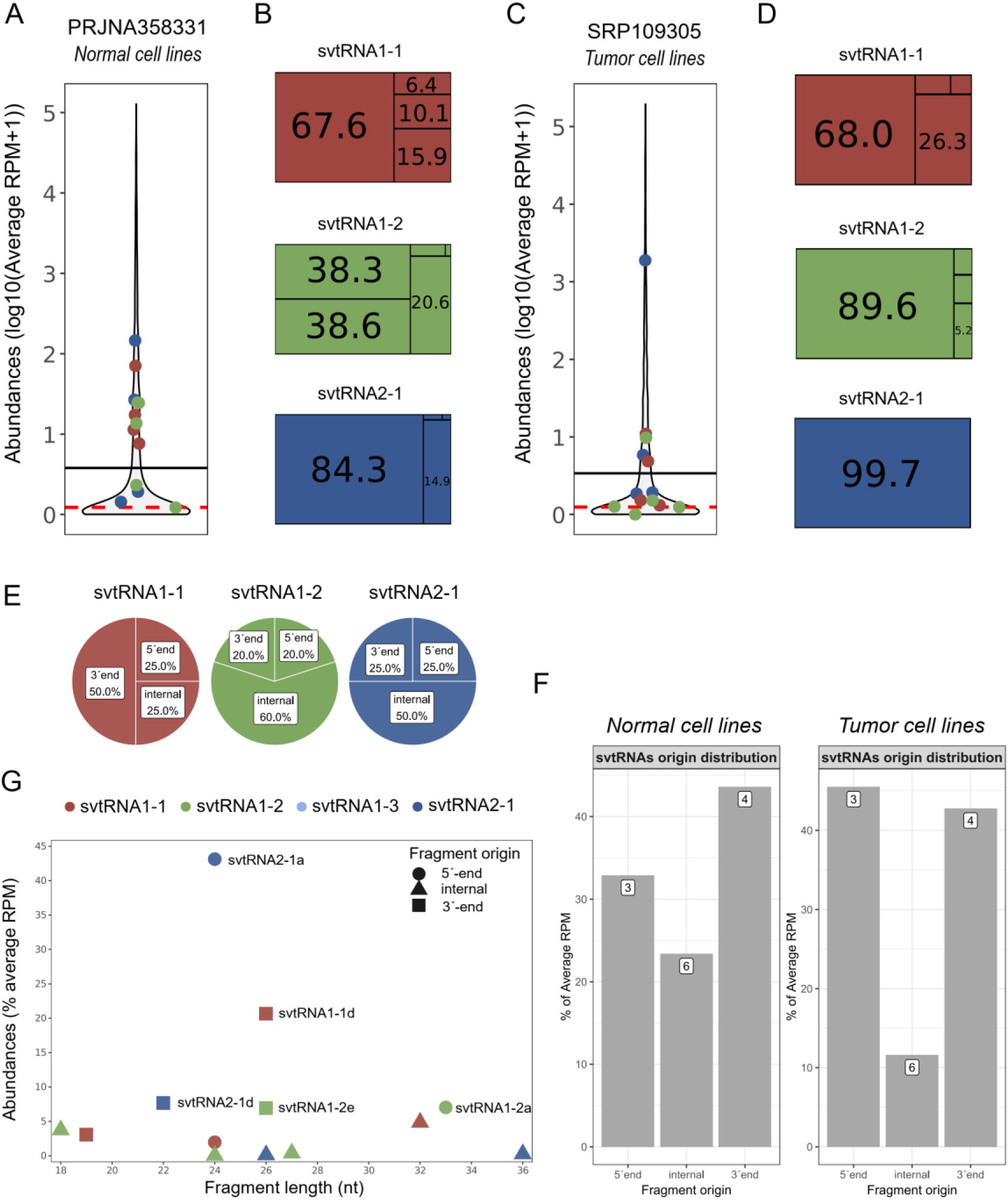
Abundances of global svtRNAs in primary and tumor cell lines datasets. **A.** Sina plot showing the abundance of global svtRNAs relative to miRNAs in primary cell lines, log10-transformed read counts are shown. Each dot represents a svtRNA in one dataset; colors correspond to the vtRNA precursor. The black line indicates the global mean of miRNA abundances, and the red line marks the median of each datatset**. B.** Treemap summarizing the relative abundance of svtRNAs within each vtRNA precursor (percentage numbers of svtRNAs with less than 5% are not shown). **C.** Sina plot showing the abundance of global svtRNAs relative to miRNAs in tumor cell lines. Each dot represents a svtRNA in one dataset; colors correspond to the vtRNA precursor. The black line indicates the global mean of miRNA abundances, and the red line marks the first quartile**. D.** Treemap summarizing the relative abundance of svtRNAs within each vtRNA precursor (percentage numbers of svtRNAs with less than 5% are not shown). **E.** Pie chart of the distribution of svtRNAs according to their positional origin within vtRNAs (5′ end, internal, or 3′ end), shown as counts. **F.** Bar plot of the distribution of svtRNAs in primary and tumor cell lines according to their positional origin within vtRNAs (5′ end, internal, or 3′ end), shown as abundance proportions. **G.** Dot plot integrating svtRNA abundance in primary cell lines, fragment length, and positional origin.

**Supplementary Figure 4.**
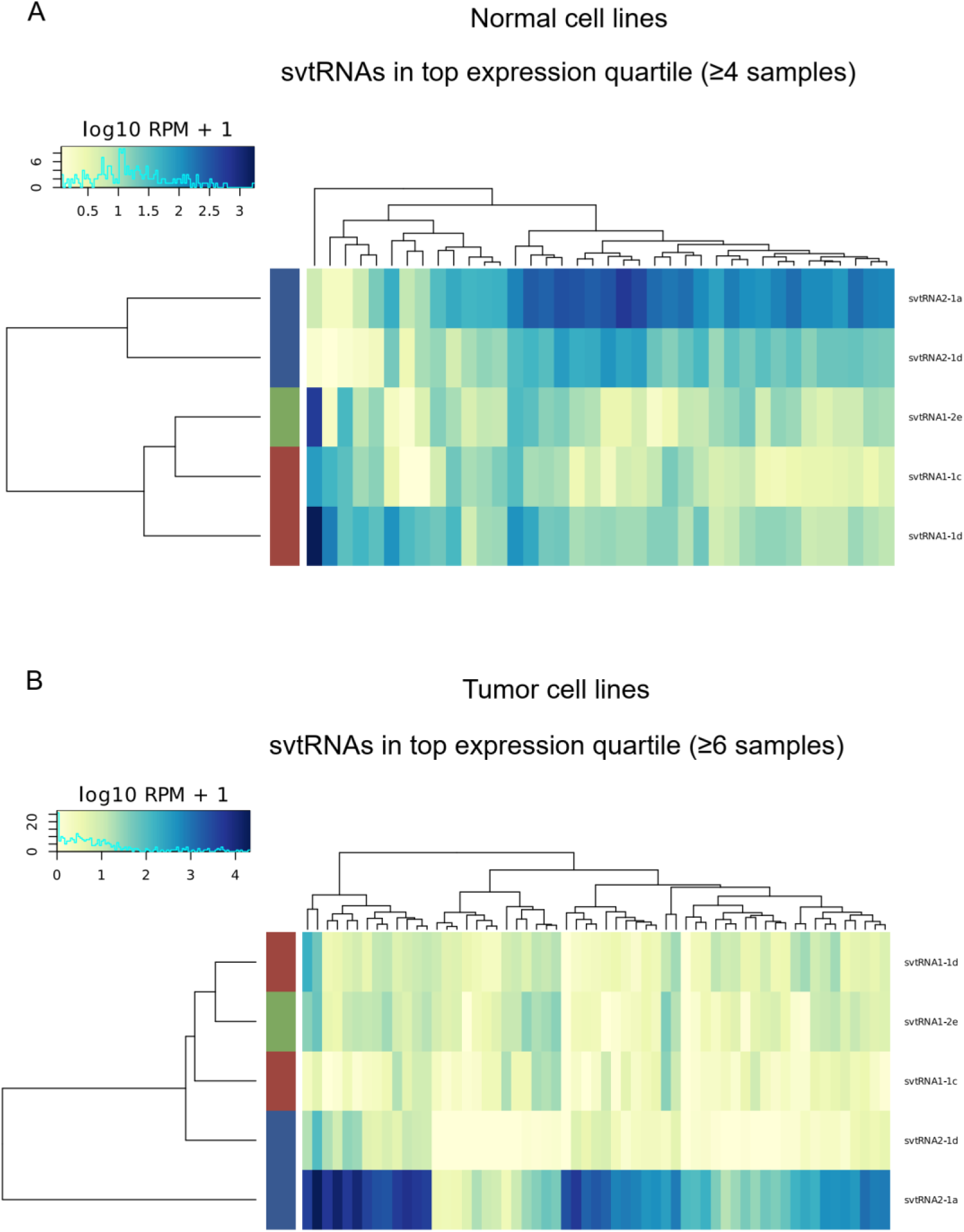
Hierarchical clustering of highly expressed Total svtRNAs in normal and tumor cell lines. Heatmaps show svtRNAs in the upper expression quartile detected in ≥4 normal samples (A) or ≥6 tumor samples (B). Values correspond to log10(RPM+1). Clustering was performed using Euclidean distance on both rows and columns. Colored side bars denote vtRNA precursor families.

**Supplementary Figure 5.**
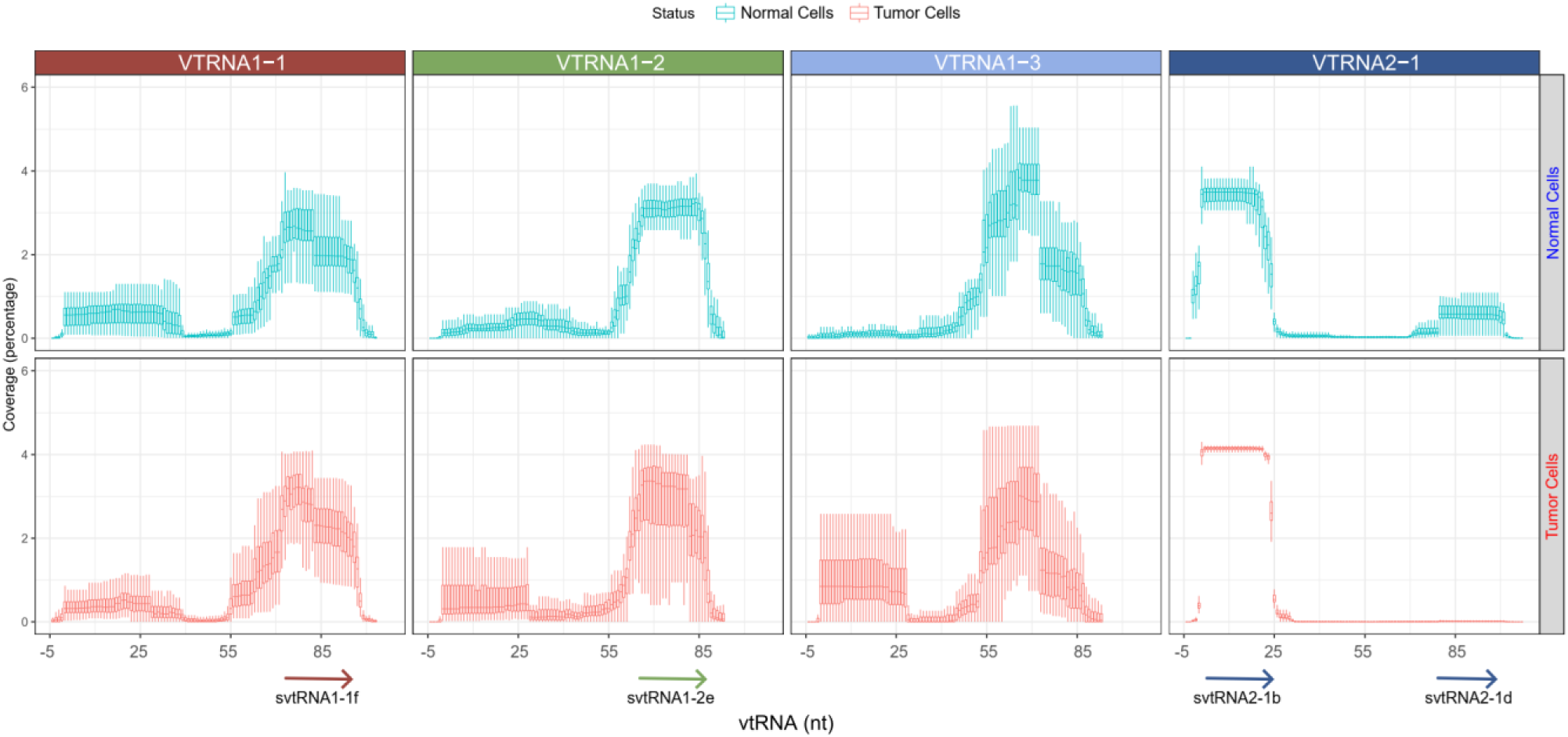
Per-base coverage profiles of vtRNAs in normal and tumor cell lines. Per-nucleotide coverage across vtRNA loci in normal primary cells (top panels) and tumor cell lines (bottom panels). Coverage values were calculated from aligned BAM files and normalized within each vtRNA. Plots show coverage distribution along vtRNA coordinates. Arrows indicate the positions of abundant miRNA-like svtRNAs identified in this study.

